# Beyond forced choice: hyper-realistic AI faces reveal a four-cluster perceptual geometry of emotion

**DOI:** 10.64898/2026.06.02.729704

**Authors:** Bliss Cui, Peter Bex

## Abstract

Forced-choice paradigms in facial emotion perception research require observers to commit to a single label and discard any simultaneous activation of related categories that participants might otherwise express. We introduce a multi-select rating paradigm in which 112 participants rated each of 52 facial-expression stimuli on 13 emotion categories using continuous 0-to-10 sliders, alongside a separate authenticity slider. Stimuli were generated using OpenAI’s DALL-E 3 from an orthogonal four-factor design (emotion, race, gender, age), allowing demographic balance and compositional uniformity more readily than photograph-based stimulus sets permit. Linear mixed-effects models with participant random intercepts and logistic regression with cluster-robust standard errors were applied to 5,824 trials. Perceived authenticity reliably predicted agreement between the participant’s top-rated category and the AI-prompted target emotion (odds ratio 1.03 per +1 unit, *p* < .001), and a principal-components analysis of the 13-emotion rating matrix recovered three interpretable dimensions accounting for over half of trial-level variance, with the first two corresponding to the canonical valence and arousal axes. Hierarchical clustering exposed a tight cluster of self-conscious negative emotions (sadness, embarrassment, shame) that does not align with the canonical basic-versus-complex emotion distinction. Collapsing the participant’s top-rated emotion to its cluster raised the trial-level prompt-agreement rate from 43.1% to 86.9% (*Δ* = +43.8 percentage points), indicating that participants’ most common departures from the prompted target fell within rather than across clusters. Among Autism-Spectrum Quotient subscales, the Social-Skill subscale showed a robust interaction with contempt-target faces (*p* < .001), with higher Social-Skill scores predicting elevated contempt ratings of approximately 1.2 points on the 0-to-10 scale. Two further secondary patterns emerged in the AQ-subscale-by-emotion grid, including an opposing-direction effect of Imagination and Attention-to-Detail subscales on awe ratings. We acknowledge that some of the recovered structure may partly reflect statistical regularities of the AI-generation pipeline rather than human perceptual geometry alone, and we discuss this caveat at length below. Multi-select rating with hyper-realistic AI-generated stimuli produces a richer perceptual probe than the forced-choice paradigms that have dominated this literature, and reveals subscale-level autistic-trait associations that AQ-total analyses obscure.

## Introduction

Decades of research on facial emotion perception have relied almost exclusively on forced-choice paradigms in which observers select a single label from a fixed set of categories, typically the six basic emotions proposed by Ekman (1992): anger, disgust, fear, happiness, sadness, and surprise. The dominance of this paradigm reflects both historical convention and methodological convenience: forced-choice produces a clean accuracy metric and supports straightforward between-group comparison (Calvo & Lundqvist, 2008; Keating & Cook, 2020). The same paradigm choice, however, discards a substantial portion of the perceptual signal that participants may provide. When the experimenter requires a single categorical answer, the simultaneous activation of related categories, the partial perception of mixed emotional content, and the perceived authenticity of the expression itself are all collapsed into the binary correct or incorrect outcome of the trial.

Recent work has begun to push beyond this constraint. Cowen and Keltner (2017) demonstrated that observers reliably distinguish 27 distinct categories of self-reported emotion in response to short videos, organized along continuous gradients rather than into mutually exclusive categories. Cordaro and colleagues (2018) reported 22 emotion categories with universal and culturally variable components in cross-cultural facial expression ratings, and Cowen and colleagues (2021) extended these findings to 16 facial expression categories occurring in similar contexts worldwide. Across this line of work, two methodological themes recur: emotion categories appear to be fuzzy and partially overlapping rather than discrete, and rich individual-difference structure exists in how observers map facial cues onto those categories. The forced-choice paradigm cannot easily capture either.

A complementary methodological development concerns stimulus design itself. Generative-AI image models now produce face images that observers cannot reliably distinguish from photographs of real people, and that are sometimes rated as more real (Nightingale & Farid, 2022; Miller, Steward, Witkower, Sutherland, Krumhuber, & Dawel, 2023). For face-perception research, this opens stimulus-design opportunities that photograph-based stimulus sets are less able to match: full factorial control over the demographic composition of the stimulus set, uniformity of pose and lighting and background across stimuli, and the elimination of model-release and identity-privacy constraints. Recent emotion-perception measures have begun to take advantage of these properties, including the PAGE assessment (Weidmann et al., 2025), which used DALL-E to construct a 20-emotion, demographically diverse test of emotion-perception ability with strong psychometric properties. The combination of fully controlled AI-generated stimuli with a multi-select rating geometry has not, to our knowledge, been previously deployed in a single design.

These same properties, however, introduce an interpretive caveat that we want to flag at the outset and return to in the Discussion. Any structure recovered from ratings of AI-generated stimuli reflects a joint product of human perceptual organization and the statistical regularities of the generation pipeline that produced the images. A model trained on internet-scale image-and-caption data may encode prototype-like configurations of facial features for each emotion label, and observers rating those images may in part be recovering that prototype geometry rather than the geometry of perception itself. We do not view this concern as fatal to the present design, since an analogous critique applies to posed photographs of trained actors, which dominate the existing literature; but the comparison between AI-generated and naturalistic stimulus sets is an empirical question that the present study cannot resolve on its own. We treat this as a methodological limitation worth foregrounding rather than relegating to a later section.

A separate literature has examined how individual differences in autistic traits relate to facial emotion perception. The Autism-Spectrum Quotient (AQ-50; Baron-Cohen, Wheelwright, Skinner, Martin, & Clubley, 2001) measures autistic traits along a continuous distribution in non-clinical populations and decomposes into five empirically validated subscales of Social Skill, Attention Switching, Attention to Detail, Communication, and Imagination (Hoekstra, Bartels, Cath, & Boomsma, 2008). Higher autistic-trait scores have been associated with differences in face-emotion perception across a range of paradigms (Murphy, Catmur, & Bird, 2024; Keating & Cook, 2020), although the interpretation of those associations remains debated. Earlier work tended to frame autistic-trait differences in emotion perception as deficits, while a more recent literature argues that the differences are better understood as different perceptual styles applied to the same stimuli (Pellicano & Heyes, 2011; Mottron, 2017). A second open question concerns the level at which autistic-trait associations operate: AQ-total scores average over five qualitatively distinct subscales, and effects detectable at the subscale level may be obscured when the total is used as a single predictor.

The present study introduces a multi-select emotion-rating paradigm that combines three methodological features: (a) participants rate every face stimulus on each of 13 emotion categories using continuous sliders, rather than choosing a single label, with a separate 14th continuous slider for perceived authenticity of the expression; (b) the 13 emotion categories include the six canonical Ekman categories alongside seven additional categories (amusement, awe, contempt, embarrassment, interest, pride, shame) chosen for their relevance to social-evaluative perception; and (c) the stimuli are fully AI-generated face images constructed via an orthogonal four-factor design (emotion, race, gender, age) that yields demographic balance and compositional uniformity across the stimulus set. We use this paradigm to ask three questions in the same dataset. First, does the multi-select rating geometry recover known structure in facial emotion perception, namely an authenticity-by-accuracy relationship and a low-dimensional latent space resembling the canonical valence-arousal circumplex (Russell, 1980; Russell & Barrett, 1999)? Second, does the rating matrix expose perceptual groupings that forced-choice paradigms would not? Third, do autistic-trait scores, measured both as the AQ-total and as the five validated subscales, moderate the perception of specific emotions in this paradigm?

## Methods

### Participants

Participants were recruited from the Northeastern University graduate and undergraduate population through the Department of Psychology subject pool (Psylink) and compensated with course credit. The final analyzed sample comprised 112 participants who provided complete data across the demographic, AQ, and emotion-rating measures. Participants ranged in age from 18 to 31 years (*M* = 19.18, *SD* = 1.75), with 90 identifying as female, 20 as male, and 2 as non-binary. The study was approved by the Northeastern Institutional Review Board (Protocol 14-09-16, “Psychophysical Study of Visual Perception and Eye Movement Control”; Principal Investigator: Peter Bex, Translational Vision Laboratory, Department of Psychology, Northeastern University); all participants provided written informed consent.

### Materials

#### Stimuli

Facial-expression stimuli consisted of 52 still images, comprising four exemplars for each of 13 target emotion categories: amusement, anger, awe, contempt, disgust, embarrassment, fear, happiness, interest, pride, sadness, shame, and surprise. The 13 categories were chosen on the basis of research-team consultation and prior work using comparable category sets for facial-expression rating (Cowen et al., 2021).

Candidate images were generated with OpenAI’s DALL-E 3 from prompts assembled via an orthogonal four-factor design varying emotion (the 13 categories above), ethnicity (the seven U.S. Census categories: White, Hispanic or Latino, Black, Asian, American Indian or Alaska Native, Native Hawaiian or Other Pacific Islander, and two or more races), gender (male, female), and age (teenager, young adult, middle-aged adult, older adult). Each prompt requested a single ultra-photorealistic, front-facing, neutral-background headshot tightly cropped to a uniform composition (forehead-top to chin-bottom), with soft even lighting, direct eye contact, and naturalistic skin texture; the verbatim prompt template appears in Supplementary Section S6. Convergent evidence supports the perceptual validity of contemporary AI-generated face images: observers fail to reliably distinguish them from photographs of real people and sometimes rate them as more real (Nightingale & Farid, 2022; Miller, Steward, Witkower, Sutherland, Krumhuber, & Dawel, 2023). We acknowledge that the present stimulus set was screened by the lead author and Principal Investigator rather than by an independent rater panel; external rating validation of the 52 selected faces is identified as a priority for replication.

From the larger candidate pool, four exemplars per emotion (52 total) were randomly drawn, balanced overall on gender (26 male and 26 female depictions across the 52 stimuli) and roughly balanced across the seven race/ethnicity categories (each category represented by 6 to 10 of the 52 stimuli) rather than weighted to U.S.-population proportions. The full demographic distribution of the final stimulus set appears in Supplementary Section S7.

#### Autism-Spectrum Quotient

The Autism-Spectrum Quotient (AQ-50; Baron-Cohen, Wheelwright, Skinner, Martin, & Clubley, 2001) is a 50-item self-report questionnaire designed to measure autistic traits in the general population. Each item is rated on a four-point Likert scale (Definitely Agree, Slightly Agree, Slightly Disagree, Definitely Disagree). Twenty-four items are positively scored (Agreement indicates an autistic trait, scoring 1 point) and 26 items are reverse-scored (Disagreement scoring 1 point), yielding a total score from 0 to 50. We also computed the five 10-item subscales established by the original publication and validated by Hoekstra, Bartels, Cath, and Boomsma (2008). Each subscale indexes a domain in which autistic individuals tend to differ from non-autistic individuals: **Social Skill** (difficulty initiating and sustaining social interaction), **Attention Switching** (difficulty disengaging attention from one focus to move to another), **Attention to Detail** (heightened orientation toward fine-grained perceptual features), **Communication** (difficulty with the pragmatic, back-and-forth aspects of conversation), and **Imagination** (preference for facts and routines over fantasy or imaginative play). Sample distribution: *M* = 18.85, *SD* = 5.93, range 8–40, median 18.

### Procedure

The study was administered online via a self-paced web application deployed on Vercel (Vercel Inc., 2024). After providing informed consent, participants completed five on-screen instruction items confirming that they would (a) use a computer rather than a phone or tablet, (b) maximize their browser window, (c) sit in a quiet place, (d) use their first instinct without rushing (with a recommended pacing of approximately 60 seconds per face), and (e) take a brief break if needed. Following instructions, participants completed all 52 rating trials. The 52 stimuli were presented one at a time in an order randomized once per session, with no image repeated within participant. On each trial the prompt read, **“Please select the emotion and authenticity level (move at least one slider above 0).”** Participants then used 13 continuous emotion sliders (one per emotion category, range 0 to 10 in 0.5-point increments, anchors labeled “least” and “most”) and one separate authenticity slider (range 0 to 10 in 0.5-point increments, anchors labeled “Inauthentic” and “Authentic”) to record their ratings. Each slider defaulted to 0 and could be left at that value if the participant did not perceive that emotion in the face. Advancing to the next trial required moving at least one emotion slider above 0. Figure 1 shows the trial interface as participants saw it.

**Figure 1.**
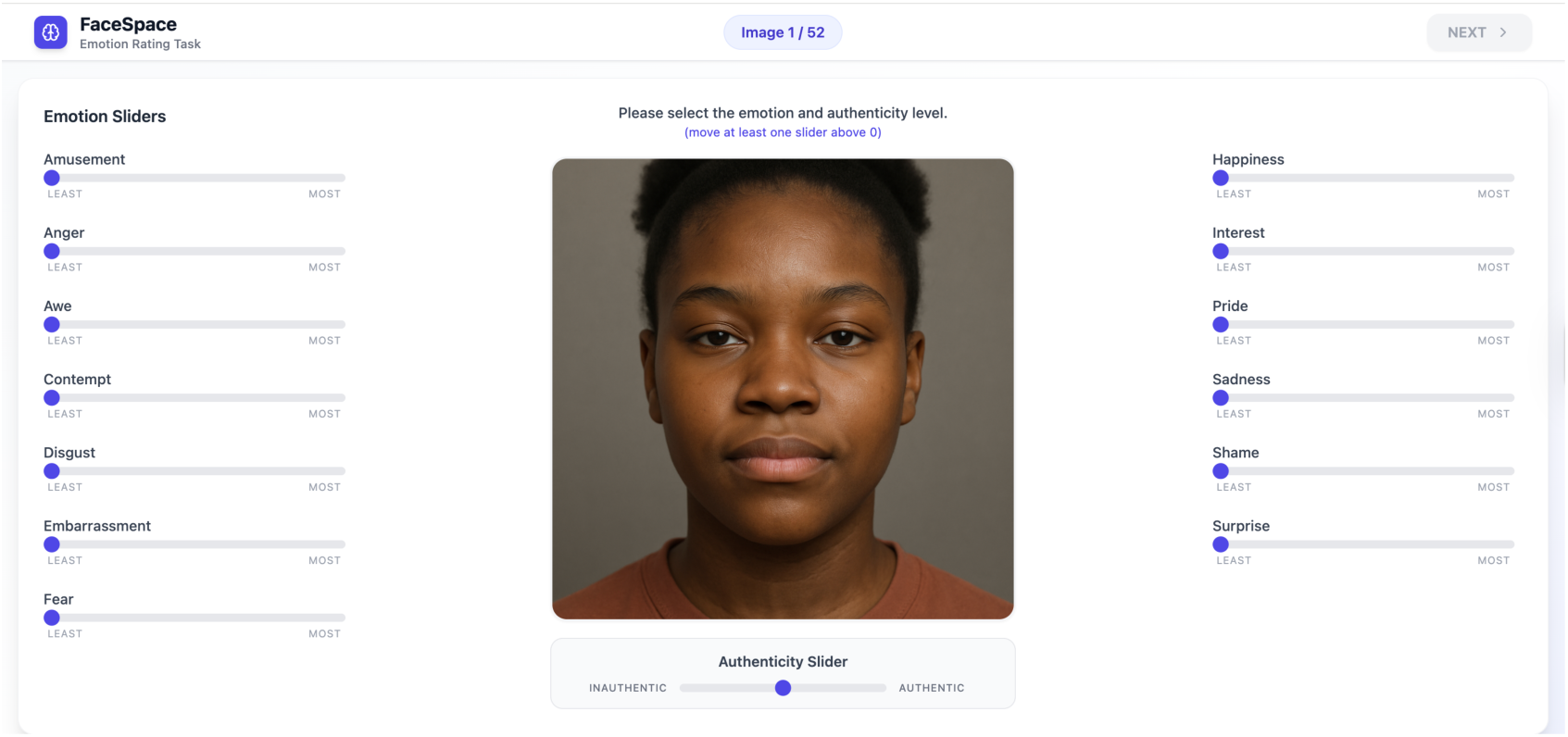
The multi-select rating interface. Trial order was randomized once per session and no image was repeated within participant. The slider state from prior trials was not visible at trial onset, so each face was rated independently. Advancing to the next trial required moving at least one emotion slider above 0; the Autism-Spectrum Quotient was administered after all 52 emotion-rating trials.

After completing all 52 rating trials, participants then completed the 50-item AQ. Administering the AQ after the rating task avoided priming participants with autism-related framing during emotion judgments. Response time per trial was recorded for descriptive transparency only and is not included as a predictor in the analyses below; the self-paced online format produced highly variable per-trial durations (median 13.6 s, with a heavy right-tailed distribution).

### Statistical Analysis

Inferential analyses respected the trial-level nesting of observations within participants. For continuous emotion-rating outcomes, we fit linear mixed-effects models with random intercepts for participants using the MixedLM procedure in statsmodels (Python; restricted maximum likelihood estimation). For binary categorization-accuracy outcomes, we used logistic regression with cluster-robust standard errors at the participant level, which provides population-average inference equivalent to a binomial generalized linear mixed model under standard assumptions. Continuous AQ scores and AQ-subscale scores were mean-centered prior to entry into models so that the intercept represents the response at the sample mean of the predictor. The effective sample for inference is the 112 participants by approximately 52 trials per participant (5,824 trials total); the mixed-effects framework draws its statistical power from this trial-level depth rather than from participant count alone, and bootstrap participant-cluster resampling (described below) was used to demonstrate robustness of headline coefficients against the participant N.

Continuous AQ was the primary autistic-trait predictor in all analyses. Three categorical operationalizations were reported as sensitivity checks: the Woodbury-Smith, Robinson, Wheelwright, and Baron-Cohen (2005) screening cutoff of ≥26 (94 below, 18 at or above the cutoff in our sample); the Baron-Cohen et al. (2001) original screening cutoff of ≥32 (109 below, 3 at or above; reported only as a descriptive comparison given the small high-AQ cell), and a sample-median split at ≥19 (61 below, 51 at or above). The categorical splits were reported in parallel rather than selected after fitting.

For analyses involving multiple comparisons within a model family, we applied the Benjamini-Hochberg false-discovery-rate correction at *q* = .05 (Benjamini & Hochberg, 1995). For the exploratory grid comparing six AQ predictors against 13 emotions (78 tests; see §3.3.2), the FDR correction was applied across the full grid rather than within rows, the more conservative framing.

Confidence intervals for headline coefficients are reported in two forms: asymptotic 95% intervals derived from the model standard errors, and participant-cluster bootstrap 95% intervals computed by resampling the 112 participants with replacement. Bootstrap iterations were 1,000 for logistic models and 200 for the linear mixed-effects models (the latter limited by computational cost).

All analyses were performed in Python 3.12 using pandas 2.1, numpy 1.26, scipy 1.11, statsmodels 0.14, and scikit-learn 1.3.

## Results

We report results in four blocks. We first establish that the multi-select rating paradigm produces internally coherent perceptual signal: perceived authenticity predicts categorization accuracy across all stimuli, the 13-emotion rating matrix has interpretable low-dimensional structure, and autistic traits, measured continuously via the Autism-Spectrum Quotient and its five validated subscales, moderate ratings of specific emotion categories. All inferential analyses use participant-level random-effects models or cluster-robust standard errors; multiple-comparison corrections use Benjamini-Hochberg false-discovery-rate adjustment within and across test families (Benjamini & Hochberg, 1995). Bootstrap confidence intervals are participant-cluster bootstraps with 1,000 (logistic models) or 200 (mixed-effects models) iterations.

### 3.1 Perceived authenticity predicts agreement with the prompted target

A note on terminology. Because the stimuli were AI-generated rather than photographs of individuals with an independently coded ground-truth emotional state, we cannot speak of categorization “accuracy” in the strict sense. Throughout, we use “accuracy” (and “correct” categorization) as a compact shorthand for agreement between the participant’s top-rated emotion category on a trial and the emotion that was specified in the DALL-E generation prompt for that stimulus. We retain the shorter term for readability in figures and model summaries but mean it in this restricted sense throughout.

Across 5,824 trials nested in 112 participants, faces that participants rated as more authentic were also faces whose prompted target emotion they more often selected as their top-rated category. We coded each trial as accurate (in the sense defined above) when the displayed expression’s prompted emotion received the highest of the 13 emotion ratings on that trial, and we modeled this binary outcome with logistic regression and cluster-robust standard errors at the participant level. Perceived authenticity, measured on a 0–10 slider in 0.5-point increments, was mean-centered (*M* = 4.32) before being included in the model, so that the intercept represents the predicted prompt-agreement at average authenticity. The slope was reliable and positive, *β* = 0.029 (*SE* = 0.007), *z* = 3.98, *p* < .001, 95% CI [0.015, 0.043]; participant-cluster bootstrap 95% CI [0.015, 0.044]. Each one-point increase in perceived authenticity multiplied the odds of correct identification by 1.03 (95% CI [1.02, 1.04]), and across the full ten-point range of the slider the odds ratio was 1.34 (95% CI [1.16, 1.55]; bootstrap 95% CI [1.16, 1.55]). The model’s McFadden pseudo-*R*² was 0.002 and Tjur’s coefficient of discrimination was 0.003. Both values are modest in absolute magnitude, as expected for a single trial-level predictor of a binary outcome with substantial stimulus-level and within-participant variation. As Figure 2 shows, the predicted probability of correct identification rose from approximately .43 at the bottom of the authenticity scale to .50 at its top, and tracked the binned empirical accuracy across the full slider.

**Figure 2.**
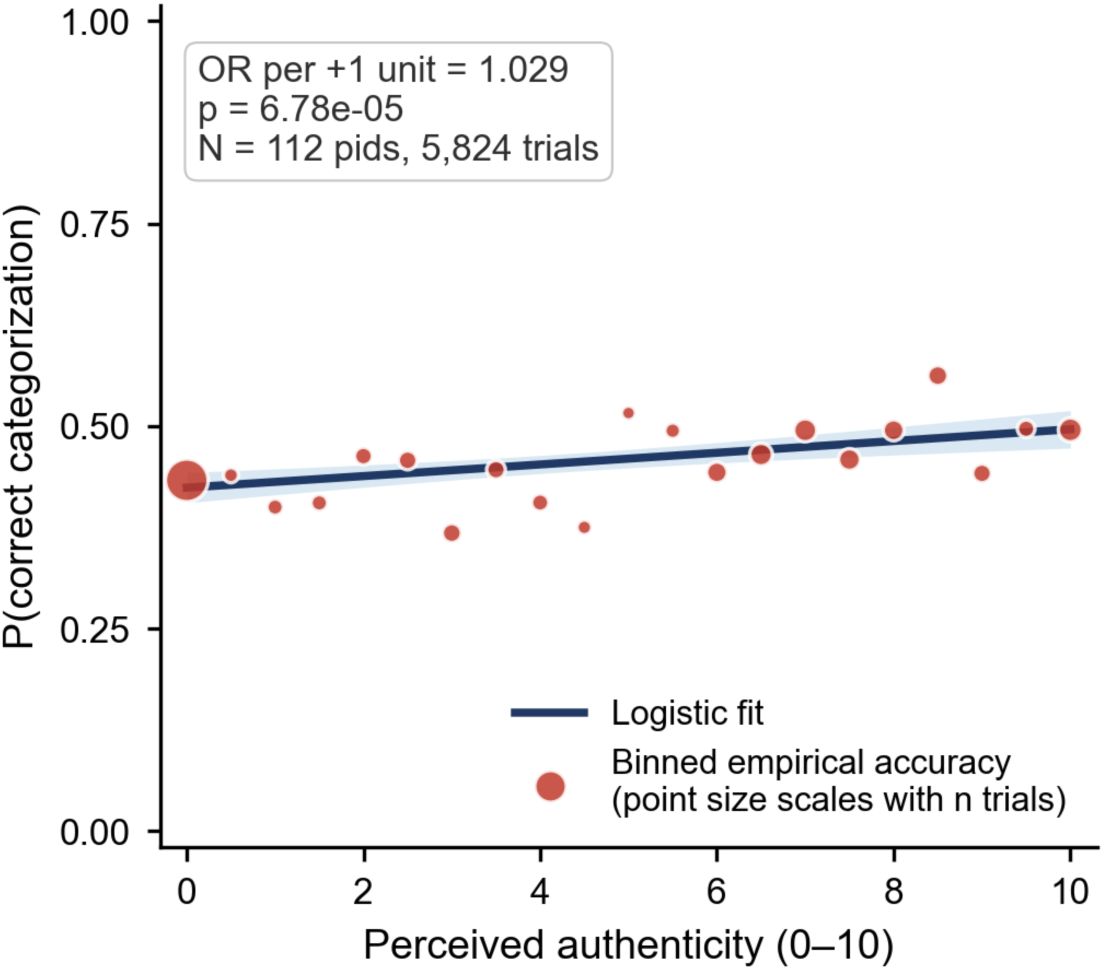
Trial-level authenticity ratings predict categorization accuracy. The blue curve is the logistic-regression fit (cluster-robust SE by participant), the shaded band is its 95% confidence interval, and red points show the empirical accuracy at each authenticity bin (point size scales with the number of trials in the bin).

The relationship was independent of autistic traits. Adding the participant’s continuous Autism-Spectrum Quotient (AQ) score as a covariate did not change the authenticity coefficient (*β* = 0.029, *p* < .001) and revealed no independent effect of AQ on accuracy (*β* = 0.004, *p* = .29). A subsequent model added the AQ × authenticity interaction term, which was likewise non-significant (*β* = 0.001, *p* = .37). The link between perceived authenticity and categorization accuracy thus appears to be a general perceptual phenomenon and was not moderated by autistic-trait individual differences in the present sample.

Overall accuracy across the 13 categories was 45.5%, above the 7.7% chance level for a 13-alternative forced-choice task but below the ceiling that prototypical-stimulus paradigms can produce. The sample-mean authenticity rating was 4.32 out of 10, with substantial spread across both the participant and stimulus dimensions (see Figure 2). The authenticity-accuracy association reported above converges with prior work on perceived authenticity of facial expressions (Calvo & Lundqvist, 2008; Murphy, Catmur, & Bird, 2024).

### 3.2 Perceptual structure of the multi-select rating matrix

Beyond validating the rating paradigm against accuracy, the trial-level rating matrix offers a window into the perceptual structure that participants impose on facial expressions when they are not constrained to choose a single label. Three complementary views (pairwise correlations, dimensionality reduction, and hierarchical clustering) converged on a coherent and partly counter-intuitive picture.

#### 3.2.1 Pairwise rating correlations

Across the 13 emotion ratings, the strongest pairwise correlations identified emotions that participants tended to perceive together when looking at the same face (Figure 3). The largest positive correlation, between amusement and happiness (*r* = +.58), reproduces the well-documented co-occurrence of these two positively-valenced emotions in continuous-rating designs (Cowen & Keltner, 2017; Cowen et al., 2021). Among the negatively-valenced emotions, embarrassment and shame (*r* = +.42), sadness and shame (*r* = +.41), and a slightly weaker pair of awe and surprise (*r* = +.30) emerged as natural perceptual clusters. The basic threat emotions of anger and disgust were correlated at *r* = +.30 (Ekman, 1992; Rozin, Lowery, Imada, & Haidt, 1999).

**Figure 3.**
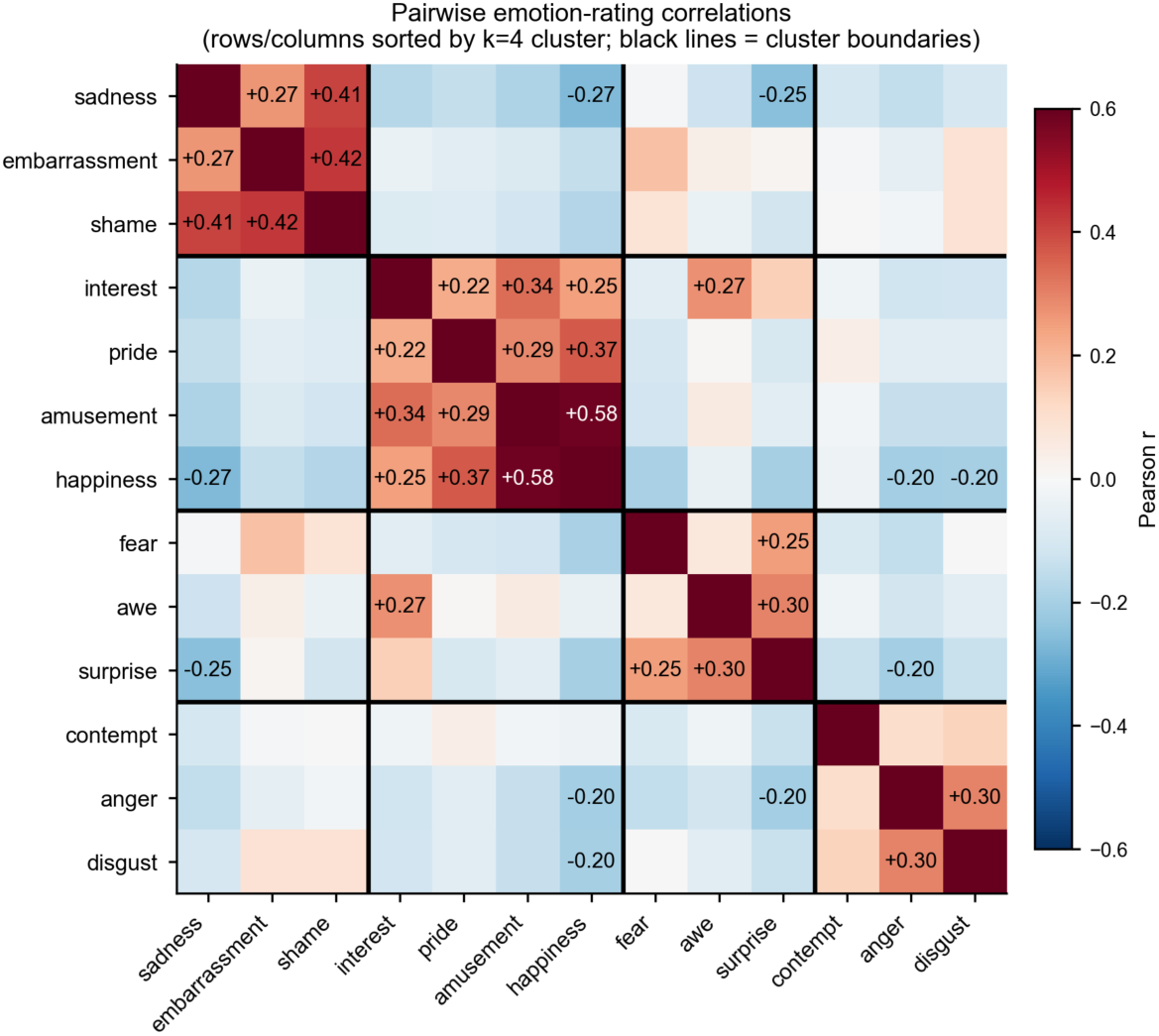
Pairwise Pearson correlations among the 13 emotion-rating columns (trial-level), reordered by Ward-linkage hierarchical clustering on the absolute-correlation distance metric (1 − |r|): if two emotions are perfectly correlated the distance is 0 and they appear adjacent on the axes. Cells with |r| ≥ 0.20 are annotated; warmer colors indicate stronger positive correlations. Black lines mark the boundaries of the four k=4 clusters reported in §3.2.3.

#### 3.2.2 Principal-components analysis

Principal-components analysis on the 5,824 × 13 rating matrix returned a low-dimensional structure that is visually consistent with the affective space that has emerged from a half-century of dimensional emotion research (Russell, 1980; Russell & Barrett, 1999). The first three components explained 21.9%, 18.5%, and 16.9% of the trial-level variance respectively, with cumulative variance of 57.3%. PC1 had its largest positive loading on happiness (+.51) and its largest negative loading on sadness (−.71), placing it close to a valence axis. PC2 was anchored at the high-arousal end by surprise (+.74) and fear (+.35) and at the low-arousal end by happiness (−.45), recovering an arousal axis. PC3 was anchored by anger (+.65) and disgust (+.47) at one pole and sadness (−.43) at the other. The biplot in Figure 4 shows the 13 emotions in PC1 × PC2 space. We emphasize that the principal components are continuous orthogonal axes that span the full 13-emotion rating space; they are not themselves the four clusters reported in §3.2.3. Each emotion has a position in PC1 × PC2 × PC3 space (a triple of loadings), and the four clusters emerge from the mutual proximity of those positions, not from any single PC.

**Figure 4.**
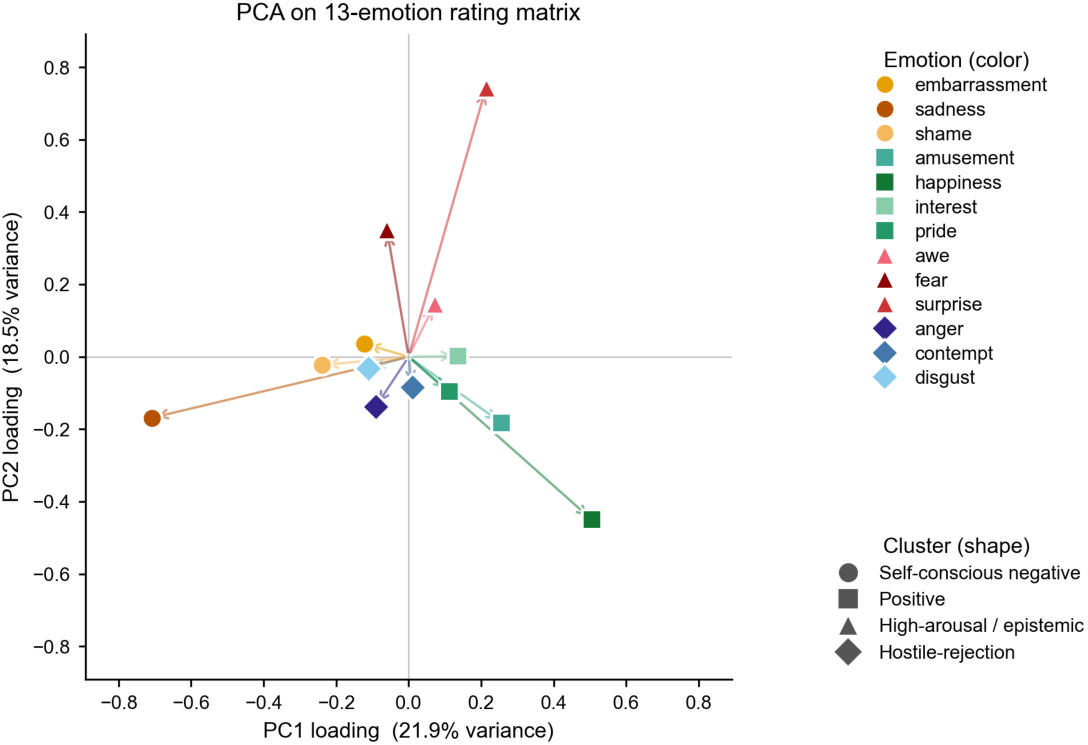
Two-dimensional PCA biplot of the 13 emotion vectors in the PC1 × PC2 plane. Arrows extend from the origin to each emotion’s projected vector tip; circles mark Ekman-canonical basic emotions and squares mark complex emotions (visual coding only; see Methods).

#### 3.2.3 Hierarchical clustering

To investigate which emotions participants perceived most similarly to one another, we applied hierarchical clustering to the 13 emotions. Hierarchical clustering builds a tree (Figure 5) by progressively merging emotions whose rating profiles are most alike, using as its similarity measure the absolute Pearson correlation between each pair of emotion-rating columns. The dendrogram yielded a clean four-cluster partition at the four-way cut, with each emotion assigned to one of four interpretable groupings.

**Figure 5.**
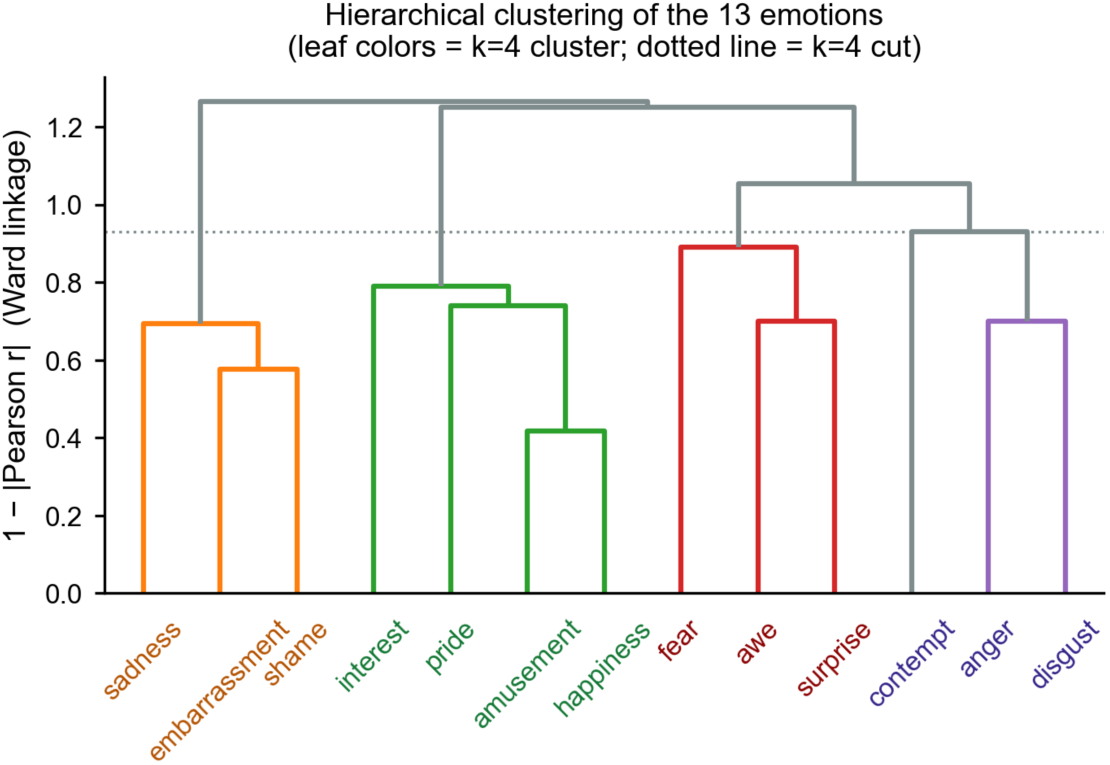
Ward-linkage hierarchical clustering of the 13 emotions on the absolute-correlation distance metric (1 − |r|). Cutting the dendrogram at four clusters yields the four interpretable groupings reported in §3.2.3.

The first cluster comprised three negatively-valenced, self-referential emotions: **sadness, embarrassment, and shame**. This grouping aligns with the self-conscious-emotion family that involves real or imagined other-evaluation (Tracy & Robins, 2004; Tangney & Dearing, 2002). The second cluster comprised four positively-valenced emotions: **amusement, happiness, interest, and pride**. These share the broad positive-affect dimension long described in self-report measures (Watson, Clark, & Tellegen, 1988) and, in the case of pride, are associated with the broad smiling display family (Tracy & Matsumoto, 2008). The third cluster grouped three high-arousal, eye-widening emotions: **awe, fear, and surprise**. These share the sensory-opening morphology (widened eyes, raised brows) that has been argued to support enhanced sensory intake during epistemic engagement (Susskind et al., 2008; Shiota, Keltner, & Mossman, 2007). The fourth cluster grouped three negatively-valenced emotions of rejection or hostility: **anger, contempt, and disgust**. This corresponds closely to the moral-emotion CAD triad described by Rozin, Lowery, Imada, and Haidt (1999) and to the hostility cluster reported by Matsumoto and Ekman (2004). Each emotion was assigned to exactly one cluster; the partition is exhaustive over the 13-emotion set. We refer to these four groupings throughout the remainder of the manuscript as the self-conscious-negative, positive, high-arousal, and hostile-rejection clusters, respectively.

The four-cluster structure recovered from rating geometry is informative for two reasons. First, it emerges without any researcher-imposed categorical structure: each cluster is the data-driven product of how participants used the 13 sliders, not a label assigned a priori. Second, the partition cuts across two of the field’s standard taxonomic distinctions in a way that points toward a hybrid rather than either-or organization. Items from the Ekman (1992) basic-emotion list are distributed across all four clusters (anger and disgust in the hostile-rejection cluster; happiness in the positive cluster; sadness in the self-conscious-negative cluster; fear and surprise in the high-arousal cluster), and self-conscious emotions in the Tracy and Robins (2004) sense (shame, embarrassment, pride) are distributed across the self-conscious-negative cluster (shame, embarrassment) and the positive cluster (pride).

The structure thus supports both views of emotion organization at once. The existence of compact, theoretically coherent clusters at the top cut of the dendrogram is consistent with a categorical organization of facial emotion perception: certain groupings of emotions recur as recognizable units in participants’ rating geometry. At the same time, the substantial spread of trial-level ratings within each cluster (and the continuous correlations among emotions across cluster boundaries reported in §3.2.1) is consistent with a dimensional, individual-differences view in which observers can differ in where their perception of any given face falls within the broader cluster structure. Context, individual perceptual style, and autistic-trait variation (§3.3) are among the factors that may modulate this within-cluster spread.

#### 3.2.4 Cluster-level prompt-agreement

The four-cluster structure recovered above suggests a complementary view of the prompt-agreement analysis in §3.1: confusions between emotions that belong to the same cluster (for example, embarrassment rated as sadness, or amusement rated as happiness) may reflect within-cluster perceptual blending rather than failure to perceive the prompted target. To quantify this, we recomputed the trial-level agreement rate after collapsing both the participant’s top-rated emotion and the prompted target to their k = 4 cluster (using the partition reported in §3.2.3). Emotion-level agreement across the 5,824 trials was 43.1% (participant-cluster bootstrap 95% CI [42.0%, 44.2%]). Cluster-level agreement, scored as a match whenever the participant’s top-rated emotion fell in the same k = 4 cluster as the prompted target, rose to 86.9% (95% CI [85.7%, 88.0%]), a +43.8-percentage-point lift. An alternative cluster-level definition that selects the cluster with the highest summed within-cluster rating (rather than the cluster of the top-1 emotion) produced an almost identical 87.7% agreement rate (95% CI [86.7%, 88.7%]). The lift was not uniform across clusters: the self-conscious-negative, high-arousal/epistemic, and hostile-rejection clusters each reached ≥93% cluster-level agreement, whereas the positive cluster (amusement, happiness, interest, pride) reached 71%, reflecting that positive-valenced ratings spread across cluster boundaries more often than the other three clusters’ ratings did. The pattern is consistent with the four-cluster geometry being the level at which participants’ rating choices most reliably track the AI-prompted target, with within-cluster substitutions accounting for the majority of the gap between emotion-level and cluster-level agreement.

### 3.3 Autistic-trait moderators of emotion perception

Both the AQ-total score and its five validated subscales (Baron-Cohen, Wheelwright, Skinner, Martin, & Clubley, 2001; Hoekstra, Bartels, Cath, & Boomsma, 2008) were entered as continuous predictors in linear mixed-effects models testing AQ × target-emotion interactions on emotion-rating intensity.

#### 3.3.1 Higher Social-Skill scores predict elevated contempt ratings to contempt-target faces

Our primary autistic-trait finding was a robust interaction between the AQ Social-Skill subscale and an indicator coding whether each trial showed a contempt-target face (i.e., whether the displayed expression was one of the four images intended to convey contempt). We modeled trial-level contempt ratings as a function of AQ Social-Skill, the contempt-target indicator, and their interaction, using a linear mixed-effects model with a random intercept for each participant. The interaction term was significant, *β* = +0.20, *SE* = 0.045, *t* = 4.51, *p* < 10⁻⁵; participant-cluster bootstrap 95% CI [0.022, 0.368]. The marginal *R*² attributable to the focal interaction was 0.004, small in absolute terms but consistent with a single subscale-emotion combination explaining variation in a single emotion rating. Translated into the rating metric: a participant scoring at the 90th percentile of the AQ Social-Skill subscale (≈6 out of 10) gave contempt ratings on contempt-target faces that were on average about 1.2 points higher (on the 0–10 scale) than a participant at the 10th percentile (≈0). The effect was directionally consistent across all four contempt-target stimuli (Figure 7), with three of four showing positive zero-order correlations (*r*s = +.20, +.15, +.19) and the fourth showing a smaller but same-sign relationship (*r* = +.10).

**Figure 6.**
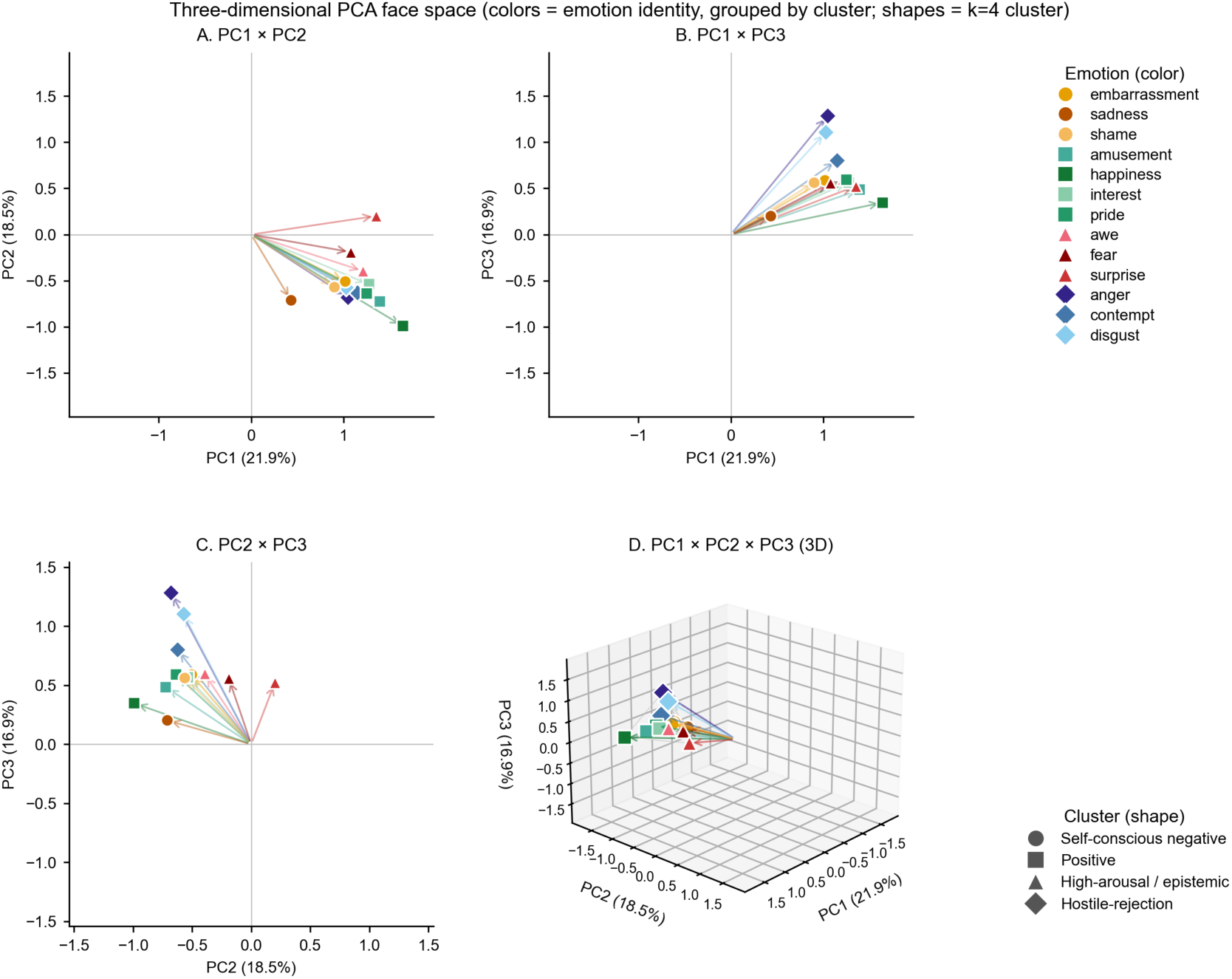
Three-dimensional PCA face space. Each of the 13 emotions occupies a single position in the PC1 × PC2 × PC3 space and is plotted as a marker at the tip of an arrow from the origin. Marker color encodes emotion identity (see right-side legend) and marker shape encodes k=4 cluster membership: ● self-conscious negative (sadness, embarrassment, shame); ▪ positive (amusement, happiness, interest, pride); ▴ high-arousal / epistemic (awe, fear, surprise); ◆ hostile-rejection (anger, contempt, disgust). Panels A–C show 2D projections of the same emotion-vector tips onto each PC pair; Panel D shows the full 3D space with convex-hull edges. The principal components are orthogonal continuous axes that span the rating space; they are not themselves the four clusters, which are formed by the mutual proximity of emotion positions across all three components.

**Figure 7.**
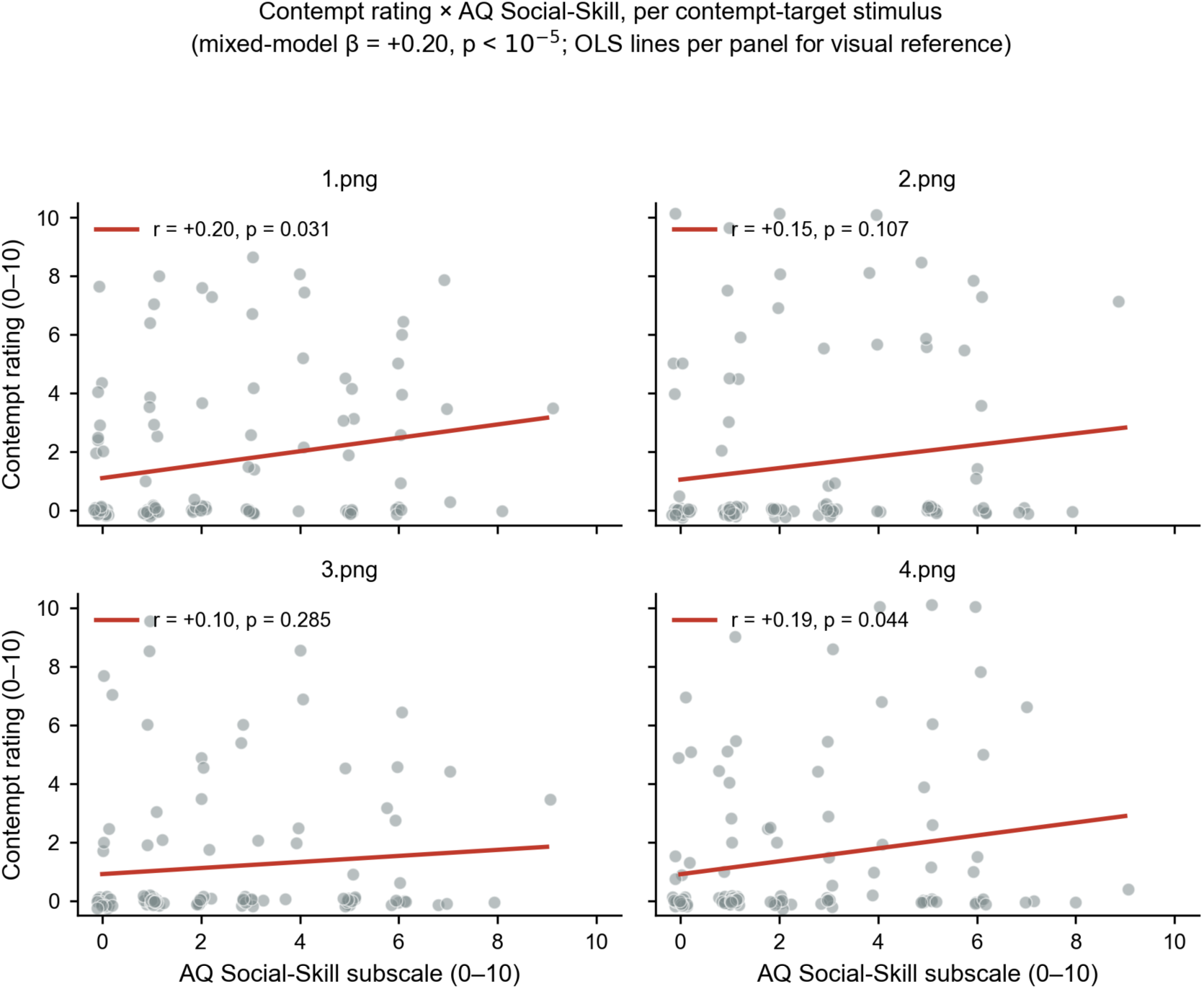
Stimulus-level robustness of the contempt × AQ Social-Skill interaction. Each of the four panels corresponds to one of the four contempt-target stimuli; within each panel, participants’ trial-level contempt ratings (jittered for visibility) are plotted against their AQ Social-Skill subscale score, with an OLS line as visual reference. This shows that the positive contempt × AQ Social-Skill association is not driven by a single stimulus: the slope is positive in three of four panels and small but same-signed in the fourth. The mixed-model β reported in §3.3.1 was estimated jointly across all four panels.

The interaction was robust across operationalization choices. It remained significant when AQ Social-Skill was discretized at the Woodbury-Smith, Robinson, Wheelwright, and Baron-Cohen (2005) clinical cutoff of 26 (*β* = +0.83, *p* = .003), at the median (*β* = +0.66, *p* = .001), and when stimulus identity was included as a fixed effect (*β* = +0.05 per AQ point, *p* = .002). Among the five AQ subscales, only Social-Skill and Communication produced significant target-modifying effects on contempt ratings; AQ Communication’s interaction was directionally consistent and somewhat smaller (*β* = +0.17, *p* = .0004), and the Attention-Switching, Attention-to-Detail, and Imagination subscales did not reach significance.

#### 3.3.2 Exploratory subscale × emotion grid

To check whether other (subscale × emotion) interactions emerged, we fit the same mixed-effects interaction model for every combination of six AQ predictors (the total score plus the five 10-item subscales) and 13 emotions, yielding 78 linear mixed-effects models. Benjamini-Hochberg false-discovery-rate correction was applied across the entire grid (*q* = .05); 12 cells survived (Figure 8). Two coherent secondary patterns emerged.

**Figure 8.**
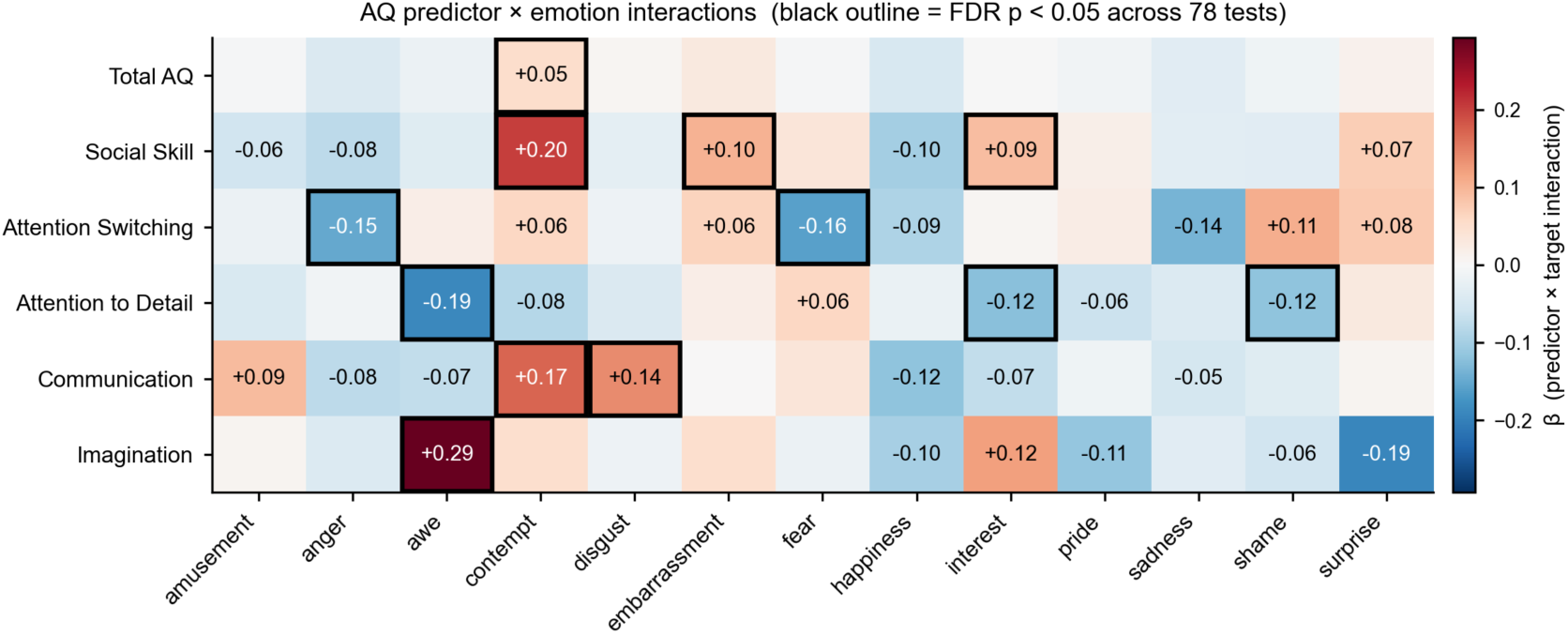
AQ-subscale × emotion interaction grid (78 LMMs total). Each cell shows the predictor × Target interaction coefficient β; cells outlined in black survive Benjamini-Hochberg FDR correction across all 78 tests at q = .05. Red shading indicates positive β (higher predictor score → elevated rating on the corresponding target-emotion trials); blue shading indicates negative β.

Awe ratings were associated with two AQ subscales in opposing directions: Imagination on awe-target trials (*β* = +0.29, *p*_FDR < 10⁻⁵) and Attention-to-Detail on awe-target trials (*β* = −0.19, *p*_FDR < .001).

AQ Social-Skill predicted higher embarrassment ratings on embarrassment-target trials (*β* = +0.10, *p*_FDR = .006) and higher interest ratings on interest-target trials (*β* = +0.09, *p*_FDR = .026). AQ Communication predicted elevated disgust ratings on disgust-target trials (*β* = +0.14, *p*_FDR = .034).

Four additional FDR-significant cells appear in Figure 8: AQ Attention-Switching predicted reduced fear (*β* = −0.16) and anger (*β* = −0.15) ratings on the corresponding target trials, and AQ Attention-to-Detail predicted reduced interest (*β* = −0.12) and shame (*β* = −0.12) ratings.

## Discussion

### 4.1 What multi-select rating reveals about facial emotion perception

The multi-select rating paradigm carried an internally coherent perceptual signal. Across all 5,824 trials, perceived authenticity predicted categorization accuracy in a way that was independent of autistic traits and survived participant-clustered standard errors at high statistical reliability. The 13-emotion rating matrix yielded a low-dimensional structure in which the first three principal components together accounted for over half of trial-level variance, with the first two components recovering the canonical valence and arousal axes long associated with the circumplex model of affect (Russell, 1980; Russell & Barrett, 1999) and a third component organized along an approach-rejection axis (anger and disgust at one pole, sadness at the other) that the two-dimensional circumplex does not predict. The need for a third interpretable axis to capture facial-expression rating variance converges with Fontaine, Scherer, Roesch, and Ellsworth (2007), who demonstrated through cross-linguistic analysis that valence and arousal are not by themselves sufficient to characterize the semantic structure of emotion concepts.

A novel finding from this rating geometry is the four-cluster structure that the rating matrix produced (Figures 5 and 6). Hierarchical clustering on the trial-level rating correlations partitioned the 13 emotions into four interpretable groupings that do not align with the canonical basic-versus-complex emotion distinction (Ekman, 1992; Cordaro et al., 2018) or with the self-conscious emotion family as conventionally defined (Tracy & Robins, 2004; Tracy & Matsumoto, 2008). Three of the four clusters group emotions sharing an appraisal pattern rather than a valence or arousal signature: the hostile-rejection cluster (anger, disgust, contempt) corresponds to the moral-emotion triad described by Rozin and colleagues (1999); the high-arousal cluster (awe, fear, surprise) groups emotions that share the eye-widening, sensory-opening morphology of epistemic engagement; and the self-conscious-negative cluster (sadness, embarrassment, shame) groups socially-evaluative distress emotions that involve real or imagined other-evaluation. The fourth cluster (amusement, happiness, interest, pride) is the positively-valenced grouping. We did not impose any of these labels in advance; the data placed each emotion into a cluster, and the cluster labels are post hoc descriptions of the resulting groupings.

Across the affective-science literature, dimensional and categorical accounts of emotion are typically presented as alternatives (Barrett, 2006; Russell, 2003; Ekman, 1992). The four-cluster structure recovered here suggests instead a hybrid organization: continuous variation along three meaningful axes, with local clustering of emotions that share appraisal content. This view treats emotion categories as attractor regions in a continuous perceptual space rather than as discrete natural kinds or as wholly constructed labels.

The attractor framing has both theoretical and empirical traction. Theoretically, it preserves the insight of categorical accounts that certain emotional configurations are recurrent and stable across observers and stimuli, while also preserving the insight of dimensional accounts that between-category variation is continuous rather than abrupt. Each of our four clusters can be read as a region of relative perceptual stability, with neighboring emotions reachable via continuous deformation through the rating geometry. This view aligns with the broader use of attractor-state frameworks in affective and cognitive science (Coan, 2010; Camras, 2011), in which complex behavioral states are characterized as low-dimensional regions of a high-dimensional dynamical system rather than as discrete states.

The attractor view makes at least one prediction that neither pure categorical nor pure dimensional accounts make. Within-cluster emotions should be more perceptually substitutable than between-cluster emotions of comparable valence, because they share an attractor basin even when their conceptual labels differ. A categorical account would predict no substitutability (natural-kind boundaries should be sharp); a pure dimensional account would predict that substitutability scales primarily with valence-arousal distance. Our cross-emotion correlations are at least consistent with the attractor prediction: within-cluster amusement and happiness co-rate at *r* = +.58, while amusement and surprise, which sit in different clusters, co-rate at *r* = −.06. Cleanly distinguishing attractor from dimensional accounts would require pairs matched on valence and arousal that fall on opposite sides of a cluster boundary, which the present 13-emotion set does not deliver. We therefore frame the attractor view as one consistent interpretation of the rating geometry rather than as a finding that the present data uniquely support.

### 4.2 What multi-select rating reveals beyond accuracy

Forced-choice paradigms reduce trial-level emotion perception to a single bit: correct or incorrect (Calvo & Lundqvist, 2008; Keating & Cook, 2020). The multi-select rating geometry preserves more information. Three examples from the present data are illustrative. First, the strongest cross-emotion rating correlations (amusement with happiness, *r* = +.58; embarrassment with shame, *r* = +.42; awe with surprise, *r* = +.30) reflect coherent perceptual blends that share appraisal content. In a forced-choice design these would be recorded as confusions or errors; here they are part of the structural signal that recovers the four-cluster geometry. Second, the trial-level authenticity slider revealed that perceived genuineness predicts prompt-agreement reliably and across all stimuli (odds ratio per +1 authenticity unit = 1.030, *p* < .001). This relationship would not be measurable in a forced-choice design that lacks a continuous authenticity dimension; even paradigms that compare posed to spontaneous expressions estimate the relationship between stimulus categories rather than within them. Third, the cluster-collapse analysis in §3.2.4 showed that emotion-level prompt-agreement (43%) nearly doubled when collapsed to the four-cluster partition (87%), revealing that the majority of participants’ “errors” were not categorical failures but within-cluster substitutions, a distinction that a forced-choice design cannot make.

The combination of a multi-select rating geometry with fully AI-generated stimuli also opens design space that prior face-perception research has not occupied. Recent work has established that contemporary generative models produce face images that observers cannot reliably distinguish from photographs of real people, and which are sometimes rated as more real (Nightingale & Farid, 2022; Miller, Steward, Witkower, Sutherland, Krumhuber, & Dawel, 2023). The present stimulus set exploits this by orthogonally crossing four demographic factors (race, gender, age, target emotion) at a level of factorial completeness that is more difficult for photograph-based stimulus sets to achieve. The PAGE emotion-perception assessment (Weidmann et al., 2025) used a similar DALL-E-based generation strategy with a 20-emotion category set; the present design pairs that strategy with a multi-select rating geometry rather than a forced-choice categorization.

The demographic balance of the resulting stimulus set is itself a methodological contribution worth highlighting. Mainstream face-perception stimulus sets such as NimStim (Tottenham et al., 2009) and KDEF (Lundqvist, Flykt, & Öhman, 1998) are built primarily from young, predominantly White adult models, and have been criticized for limiting the generalizability of face-perception findings beyond demographically narrow samples (Strohminger, Gray, Chituc, Heffner, Schein, & Heagins, 2016). Building a photograph-based set with the same demographic balance we achieved here (seven race or ethnicity categories crossed with two genders and four age categories, all within each of 13 emotion categories) would require a recruitment, photography, and consent pipeline that is logistically substantial. Generative AI offers a different path: any combination of factors can be sampled to arbitrary balance at no extra cost. The resulting design covers a broader region of the demographic space than a typical face-perception study and reduces the risk that emotion-perception conclusions are silently calibrated to a narrow demographic distribution.

### 4.3 Higher AQ Social-Skill scores predict elevated contempt ratings

The most robust autistic-trait finding was an interaction between the AQ Social-Skill subscale and the contempt-target indicator. Higher Social-Skill scores (indexing greater self-reported difficulty with social interaction) were associated with elevated contempt ratings on contempt-target faces. The effect size in the rating metric was substantial for an individual-difference effect: participants near the top of the Social-Skill distribution gave contempt ratings on contempt-target faces approximately 1.2 points higher (on the 0 to 10 scale) than participants near the bottom. The interaction was robust across operationalizations, including the continuous Social-Skill score, two categorical splits of the AQ-total, and a model controlling for stimulus identity, and across the four contempt-target images.

Two possible explanations of this pattern are worth considering. The first locates the effect in a more general literature on hostile-attribution tendencies, in which individuals who report greater difficulty with social interaction also show evaluative biases toward attributing negative social intent to ambiguous social cues (Crick & Dodge, 1994). The second locates the effect in research suggesting that autistic individuals may apply differently calibrated rather than less accurate social-evaluative templates to face stimuli (Klin et al., 2002; Keating & Cook, 2020), consistent with broader calls to reframe autistic-trait differences in social-cognitive performance as different processing styles rather than as deficits (Pellicano & Heyes, 2011; Mottron, 2017). Distinguishing between these interpretations would require a paradigm that varied the ambiguity of the contempt cue, the available alternative interpretations, or both.

### 4.4 An awe paradox and a Social-Skill-by-negative-emotion cluster

Two further patterns emerged in the exploratory subscale-by-emotion grid. The first is an opposing-direction pair of effects on awe ratings: higher AQ Imagination scores predicted elevated awe ratings on awe-target faces, while higher AQ Attention-to-Detail scores predicted reduced awe ratings on the same faces. Awe is theorized to involve appraisals of perceptual or conceptual vastness paired with a need for cognitive accommodation (Keltner & Haidt, 2003; Shiota, Keltner, & Mossman, 2007), and an imaginative cognitive style may facilitate the accommodation step while a strongly detail-oriented style may interfere with the holistic vastness percept. This finding is single-emotion, and replication with new stimuli would be needed before any strong claim can be made.

The second pattern is broader. The two AQ subscales most directly tied to interpersonal functioning, Social-Skill and Communication, both predicted elevated ratings of negatively-valenced and socially-evaluative emotions on their target trials: contempt for both subscales, embarrassment and interest for Social-Skill, and disgust for Communication. By contrast, the cognitive-style subscales (Attention-Switching, Attention-to-Detail, and Imagination) did not produce the same pattern. The convergence across multiple emotions suggests that interpersonal-functioning AQ subscales modulate ratings of emotions whose appraisal content centers on social evaluation, while the cognitive-style subscales modulate different aspects of the rating process.

### 4.5 AI-generated stimuli: what the present data can and cannot tell us

We flagged at the end of the Introduction a methodological caveat that warrants a sustained discussion here, because it bears on the interpretation of every result above. The four-cluster perceptual geometry, the within-cluster substitutability documented in §3.2.4, and the AQ-subscale-by-emotion interactions in §3.3 were all recovered from observers’ ratings of AI-generated faces. Any such result is a joint product of two factors: (a) the perceptual organization that human observers bring to facial emotion categorization, and (b) the statistical regularities that the generative model encoded for each prompted emotion label during its training. A reviewer can legitimately ask which of these two sources contributes more to the structure we report. We cannot definitively rule out the possibility that the four-cluster partition reflects, at least in part, the way DALL-E 3 has internalized facial-feature prototypes for each of our 13 emotion labels, rather than only the way humans organize facial-emotion perception.

Three considerations qualify this concern but do not eliminate it. First, an analogous critique applies to the dominant photograph-based stimulus sets used in this literature. Posed expressions produced by trained actors (the basis of widely used sets such as NimStim, KDEF, and the Ekman & Friesen (1976) Pictures of Facial Affect) reflect actors’ learned production schemata for prototypical expressions of each emotion category, which are themselves shaped by cultural conventions, director instructions, and the historical emphasis on the six Ekman categories. Findings recovered from such sets are, in the same sense, joint products of human perceptual organization and a production pipeline whose biases are at least as opaque as those of a generative model (see Calvo & Lundqvist, 2008, and Krumhuber, Skora, Küster, & Fou, 2017, on the limits of posed-expression validity). The relevant comparison is therefore not between AI-generated stimuli and “the truth,” but between two pipelines (posed-photograph and generative), each with its own stylization. Whether they recover the same perceptual structure when matched for category set and rating geometry is an empirical question worth addressing directly in future work.

Second, the multi-select rating geometry itself attenuates a strong version of the artifact concern. If the four clusters were primarily an imprint of the generation pipeline, participants would still need to use their 13 sliders in a way that mirrors that imprint; the rating pattern is supplied by participants, not by the model. The within-cluster correlations (*r* between .30 and .58 for the strongest pairs), the across-participant consistency of the partition, and the AQ-subscale interactions that vary across individuals all require that observers be doing something systematic with the rating sliders. A pure stimulus-property account would not by itself predict that, for example, AQ Social-Skill scores would moderate contempt-target ratings differently from amusement-target ratings.

Third, the field has begun to develop methods for partly disentangling stimulus-side from perception-side contributions that are worth highlighting as natural follow-ups to the present design. Participant-driven face-synthesis approaches, in which a genetic algorithm or comparable evolutionary procedure is used to construct individualized exemplars of specific affects from each participant’s own perceptual responses (Binetti, Roubtsova, Carlisi, Cosker, Viding, & Mareschal, 2022; Jack & Schyns, 2017), and reverse-correlation procedures that recover the perceptual templates observers apply to face images (Dotsch, Wigboldus, Langner, & van Knippenberg, 2008; Adolphs, Nummenmaa, Todorov, & Haxby, 2016), remove the experimenter (and the generative model’s training data) from the loop and produce stimuli whose structure can be attributed more directly to the participant’s perceptual geometry. The present paradigm is complementary to these approaches rather than competitive with them: the AI-generated set affords full demographic factorial design at scale, which participant-driven approaches do not, while participant-driven approaches afford a cleaner attribution of recovered structure to perception, which the present design does not [TODO: cite Bliss’s preprints here]. Cross-validating a four-cluster geometry across AI-generated stimuli, posed photographs, and participant-driven synthesis would constitute strong evidence that the geometry indexes human perceptual organization rather than the regularities of any single stimulus pipeline.

We therefore foreground this caveat without retreating from the substantive claims of the paper. The four-cluster structure, the authenticity-agreement relationship, and the AQ-subscale interactions are findings about how human observers used a multi-select rating instrument applied to a particular family of stimuli. Their generalization to naturalistic-photograph and dynamic-expression settings is an empirical question that the design described here was not built to answer alone. Three considerations informed our judgment that the design was nevertheless worth running: the demographic balance afforded by AI-generated stimuli is a genuine advance over posed-photograph sets, the within-stimulus-set findings are interpretable on their own terms even if generalization is partial, and the comparison between stimulus pipelines is a question the field needs to address regardless of which side of it any individual study sits on.

### 4.6 Other limitations and future directions

Beyond the stimulus-pipeline caveat addressed in §4.5, several further limitations qualify the conclusions above. The participant sample was drawn from a single undergraduate population (mean age 19.2 years, predominantly female respondents), so generalization across age cohorts and gender distributions is untested. The default-zero rating design conflates “not present” with “not yet considered,” a deliberate choice to avoid biasing participants by forcing every slider above zero but one that a future design variant could disentangle. The present autistic-trait conclusions are based on AQ scores in a non-clinical undergraduate sample rather than on a clinical autism diagnosis, and we did not include a measure of alexithymia, which has been argued to confound autistic-trait associations with emotion-perception measures (Bird & Cook, 2013; Murphy, Catmur, & Bird, 2024). Finally, AI-generated faces are known to tend toward more statistically average configurations than photographs of specific individuals (Dunn et al., 2024); because the present design targets the perception of emotion categories rather than identity recognition or distinctive-face memory, this caveat is less consequential than it would be for face-recognition paradigms, and prompts can in principle be engineered to include atypical or distinctive features, but it should be acknowledged when generalizing to naturalistic-photograph settings.

Productive next steps include replication in clinically diagnosed autism samples and in age cohorts outside the undergraduate range, alexithymia-controlled designs, and the extension of the multi-select paradigm to dynamic video stimuli (Cordaro et al., 2018; Cowen et al., 2021). The full subscale-by-emotion grid in Figure 8 also identifies several leads (for example, AQ Attention-Switching with reduced fear and anger ratings) that the present study did not interpret in depth and that warrant targeted follow-up.

Three translational directions are worth noting. First, dimensional emotion-perception assessment in alexithymia research. Alexithymia is robustly associated with deficits in labeling static facial expressions (Bird & Cook, 2013), and a recent systematic review and meta-analysis (Willis, Miller, More, & de la Piedad Garcia, 2025) of 24 studies estimated a medium negative alexithymia-recognition association (*r* ≈ −.24) that was specific to static-image stimuli (all included studies used forced-choice labeling tasks by inclusion criterion, so the moderating effect concerned dynamic vs. static stimuli rather than paradigm type per se). Conspicuously absent from this literature is a measurement instrument that captures the rating-pattern signatures (reduced cross-emotion blending, compressed perceptual manifolds, atypical cluster boundaries) that have been hypothesized to characterize alexithymic emotion perception but cannot be quantified through forced-choice accuracy alone. Multi-select continuous rating provides exactly this per-participant geometric fingerprint and would be a natural substrate for an alexithymia-targeted assessment protocol.

Second, personalized stimulus generation for graded exposure therapy. Recent work in social anxiety disorder has used digitally modified therapist facial expressions to deliver intensity-titrated exposure during in-session CBT (e.g., Horigome, Yoshida, Tanikawa, Mimura, & Kishimoto, 2023), and augmented-reality systems driven by AI-generated synthetic humans are emerging in PTSD and anxiety-disorder care contexts (Javanbakht, Hinchey, Gorski, Ballard, Ritchie, & Amirsadri, 2024). A persistent constraint in these settings is the difficulty of building demographically matched, intensity-graded stimulus sequences for individual patients. Our orthogonal four-factor stimulus pipeline (race × gender × age × emotion category) addresses this directly: any combination of factors can be generated to match a patient’s salient demographic context, and graded morphs along specific emotion or demographic dimensions become low-cost to produce. The present methodology supplies a generation infrastructure that the clinical exposure-therapy literature has begun calling for.

Third, dimensional screening for hostile-attribution tendencies. The contempt × AQ Social-Skill interaction reported in §3.3.1 parallels patterns described in the hostile-attribution-bias literature (Crick & Dodge, 1994), in which individuals reporting greater interpersonal difficulty also show evaluative biases toward hostile interpretations of ambiguous social cues. A multi-select rating signature that elevates contempt-target faces, interpretable at the individual level rather than only the group level, could in principle serve as a complementary screening measure alongside existing self-report instruments. We frame these three directions as foundational targets for replication-driven clinical translation rather than as claims that the present data themselves support clinical use. Each would require validation in clinical samples, an alexithymia covariate (Bird & Cook, 2013; Murphy, Catmur, & Bird, 2024), and benchmarking against established instruments before any clinical deployment.

A more open-ended direction concerns the geometry of the rating manifold itself. The four-cluster structure recovered here invites a research program in which the shape of the manifold (cluster compactness, between-cluster separation, the relative position and elongation of the clusters in PC space) becomes the unit of analysis for individual or group differences. Such an approach would treat the manifold geometry as a phenotype, asking whether perceptual organization differs in groups defined by alexithymia (Taylor, Bagby, & Parker, 1997), depression (Leppanen, 2006), threat-related attentional bias (Bar-Haim, Lamy, Pergamin, Bakermans-Kranenburg, & van IJzendoorn, 2007), or developmental stage (Widen & Russell, 2008). The twenty-seven-emotion semantic space of Cowen and Keltner (2017) and the twelve-point circumplex of Yik, Russell, and Steiger (2011) provide comparison structures against which the present four-cluster geometry could be tested. The methodology demonstrated here would scale naturally to such comparisons.

### 4.7 Conclusion

Multi-select emotion rating, paired with AI-generated facial-expression stimuli, produces a richer perceptual signal than the forced-choice paradigms that dominate the field. The 13-emotion rating geometry recovered both the canonical valence and arousal dimensions and a third interpretable component, and partitioned the 13 emotions into four interpretable perceptual clusters (self-conscious-negative, positive, high-arousal, hostile-rejection) that do not align with the canonical basic-versus-complex distinction. Within this paradigm, autistic-trait individual differences moderate the rating of specific emotions in specific ways, with the AQ Social-Skill subscale predicting elevated contempt ratings on contempt-target faces and a broader cluster of interpersonal-functioning subscales modulating ratings of emotions with social-evaluative content. These specific subscale-by-emotion patterns are not visible when AQ-total is used as a single predictor, supporting the case for analyzing autistic-trait individual differences at the subscale level and for treating autistic-trait variation as a multifaceted axis of perceptual style rather than a unitary deficit dimension.

## Supplementary Materials

The following materials accompany the main text. Numerical tables and secondary analyses that would otherwise crowd the main text are reported here for reference and reproducibility.

### S1. Detailed sample characteristics

First-language reporting in the canonical sample of 112 participants was as follows: English, 92 participants; Chinese, 7; Spanish, 7; Korean, 3; Turkish, 1; French, 1; Gujarati, 1. The sample distribution of the Autism-Spectrum Quotient subscales is summarized in Supplementary Table S1.

**Supplementary Table S1.** AQ-50 subscale and total-score distributions in the canonical sample (N = 112). Each subscale is scored on a 0 to 10 range; the AQ Total ranges 0 to 50.

| Subscale | M | SD | Median | Range |
| --- | --- | --- | --- | --- |
| Social Skill | 2.44 | 2.25 | 2.0 | 0–9 |
| Attention Switching | 5.41 | 2.03 | 6.0 | 0–10 |
| Attention to Detail | 6.15 | 1.98 | 6.0 | 1–10 |
| Communication | 2.64 | 2.10 | 2.0 | 0–10 |
| Imagination | 2.21 | 1.42 | 2.0 | 0–6 |
| AQ Total | 18.85 | 5.93 | 18.0 | 8–40 |

### S2. AQ-categorical sensitivity analyses

Supplementary Table S2 reports the contempt by target interaction estimated under each of the categorical and continuous operationalizations of the AQ score. The model in each row is contempt ∼ AQ × target + (1 | participant), and the focal coefficient is the AQ × target interaction term. Cutoff 32 is reported only as a descriptive comparison given the small high-AQ cell (n = 3); the other operationalizations all yield a positive and significant interaction.

**Supplementary Table S2.** Sensitivity of the contempt × AQ-Social-Skill interaction across operationalization choices. The continuous-AQ Social-Skill subscale model is the same model reported in §3.3.1 of the main text.

| Operationalization | $\beta$ (AQ × target) | p | Notes |
| --- | --- | --- | --- |
| Continuous AQ Total | +0.051 | 0.0027 | primary continuous predictor |
| Continuous AQ Social-Skill | +0.203 | <.0001 | headline subscale finding |
| Continuous AQ Communication | +0.170 | 0.0004 | second positive subscale |
| Continuous AQ Attention-Switching | +0.061 | 0.2191 |  |
| Continuous AQ Attention-to-Detail | -0.082 | 0.1061 |  |
| Continuous AQ Imagination | +0.050 | 0.4872 |  |
| Cutoff 26 (Woodbury-Smith) | +0.828 | 0.0025 | 94 vs 18 |
| Cutoff 32 (Baron-Cohen) | +0.346 | 0.5783 | 109 vs 3 (underpowered) |
| Median split at 19 | +0.656 | 0.0012 | 61 vs 51 |
| Continuous + stimulus fixed effects | +0.051 | 0.0015 | stimulus-controlled |

### S3. Bootstrap confidence-interval tables

Bootstrap confidence intervals were computed by resampling participants with replacement (1,000 iterations for logistic models; 200 iterations for the linear mixed-effects model). Supplementary Table S3 reports the 2.5th, 50th, and 97.5th percentiles of the resampled coefficient distributions for the three headline focal terms in the manuscript, alongside the asymptotic point estimate and standard error.

**Supplementary Table S3.** Participant-cluster bootstrap distributions for the three focal coefficients reported in §3.1, §3.3.1, and the AQ-total overall accuracy comparison. The asymptotic point estimate (β) and standard error (SE) appear alongside the 2.5th, 50th, and 97.5th percentiles of the bootstrap distribution. All three focal terms have bootstrap 95% intervals that exclude zero, except the AQ-total overall-accuracy coefficient (consistent with the non-significant association reported in the main text).

| Coefficient | $\beta$ (point) | SE | Boot 2.5% | Boot 50% | Boot 97.5% |
| --- | --- | --- | --- | --- | --- |
| Authenticity → accuracy (logistic) | +0.0291 | 0.0072 | +0.0146 | +0.0288 | +0.0437 |
| AQ-total → overall accuracy (logistic) | +0.0038 | 0.0045 | -0.0044 | +0.0034 | +0.0114 |
| Contempt × AQ-Social (LMM, 200 boots) | +0.2029 | 0.0450 | +0.0220 | +0.1841 | +0.3680 |

### S4. Full AQ-subscale × emotion grid

Supplementary Table S4 reports the numerical version of the heatmap shown in Figure 8: each cell is the AQ predictor × Target interaction term from a linear mixed-effects model fitted to the trial-level rating of that emotion, with a random intercept for participants. β is the AQ × target interaction coefficient (in rating-scale units per AQ-subscale point), and p_FDR is the Benjamini-Hochberg false-discovery-rate-adjusted p value across the full 6 × 13 grid (78 tests). Cells in bold survive p_FDR < .05.

**Supplementary Table S4.** Full numerical results of the AQ-subscale × emotion interaction grid from Figure 8. Each cell shows the LMM AQ × target interaction β (top, rating-scale units per subscale point) and the FDR-adjusted p value (bottom, italic). Yellow shading + bold β indicates p_FDR < .05 across the full 78-test grid.

| Emotion | AQ Total | Social Skill | Attn-Switch | Attn-Detail | Communic. | Imagination |
| --- | --- | --- | --- | --- | --- | --- |
| amusement | -0.00 .890 | -0.06 .289 | -0.02 .844 | -0.04 .535 | +0.09 .069 | +0.01 .910 |
| anger | -0.04 .081 | -0.08 .276 | **-0.15 .031** | -0.01 .890 | -0.08 .309 | -0.04 .788 |
| awe | -0.02 .462 | -0.04 .572 | +0.02 .822 | **-0.19 <.001** | -0.07 .168 | **+0.29 <.001** |
| contempt | **+0.05 .026** | **+0.20 <.001** | +0.06 .462 | -0.08 .285 | **+0.17 .005** | +0.05 .727 |
| disgust | +0.01 .865 | -0.03 .772 | -0.02 .865 | -0.04 .638 | **+0.14 .034** | -0.02 .890 |
| embarrassment | +0.03 .069 | **+0.10 .006** | +0.06 .166 | +0.02 .772 | +0.00 .956 | +0.05 .572 |
| fear | -0.00 .890 | +0.04 .616 | **-0.16 .003** | +0.06 .367 | +0.03 .616 | -0.02 .865 |
| happiness | -0.05 .088 | -0.10 .190 | -0.09 .336 | -0.02 .865 | -0.12 .146 | -0.10 .527 |
| interest | -0.00 .890 | **+0.09 .026** | +0.00 .910 | **-0.12 .005** | -0.07 .102 | +0.12 .067 |
| pride | -0.01 .616 | +0.02 .772 | +0.02 .727 | -0.06 .168 | -0.01 .860 | -0.11 .067 |
| sadness | -0.03 .431 | -0.03 .822 | -0.14 .190 | -0.04 .788 | -0.05 .711 | -0.03 .865 |
| shame | -0.01 .616 | -0.03 .616 | +0.11 .067 | **-0.12 .049** | -0.04 .616 | -0.06 .616 |
| surprise | +0.01 .772 | +0.07 .462 | +0.08 .475 | +0.03 .838 | +0.01 .910 | -0.19 .134 |

### S5. Per-stimulus item statistics

For each of the 52 stimuli, we computed the proportion of trials on which the participant’s highest-rated emotion matched the displayed target emotion (accuracy), the mean rating on the target-emotion slider, the mean rating across the 12 non-target emotion sliders (off-target mean), the mean authenticity rating, and the median per-trial response time. Aggregate summaries across the four exemplars per emotion are shown in Supplementary Table S5; full per-stimulus results are available in the deposited data (claude_analysis/04_exploratory/output/m05b_per_stimulus_stats.csv).

**Supplementary Table S5.** Per-emotion summary of stimulus-level performance, averaged across the four exemplars per emotion category. Sorted by descending accuracy. Off-target ratings are averages across the 12 non-target emotion sliders on the same trial; values close to 0 indicate clear target perception, while higher off-target means indicate perceptual blending.

| Target emotion | Mean accuracy | M target rating | M off-target | M authenticity | Median RT (s) |
| --- | --- | --- | --- | --- | --- |
| anger | 91.7% | 7.55 | 0.33 | 4.41 | 13.49 |
| happiness | 90.6% | 7.34 | 0.48 | 5.33 | 11.90 |
| surprise | 87.7% | 7.44 | 0.45 | 3.37 | 13.76 |
| sadness | 85.9% | 7.12 | 0.41 | 5.26 | 13.93 |
| fear | 76.8% | 7.14 | 0.65 | 4.13 | 14.89 |
| disgust | 63.2% | 6.11 | 0.64 | 3.69 | 13.59 |
| amusement | 23.4% | 4.21 | 0.97 | 5.57 | 12.72 |
| interest | 19.4% | 1.45 | 0.63 | 4.28 | 15.47 |
| pride | 17.2% | 1.36 | 0.54 | 4.81 | 13.81 |
| shame | 10.5% | 2.33 | 0.83 | 4.69 | 13.46 |
| awe | 10.0% | 1.75 | 0.92 | 3.63 | 13.36 |
| contempt | 9.8% | 1.45 | 0.92 | 3.29 | 14.08 |
| embarrassment | 5.1% | 1.42 | 0.93 | 3.65 | 14.19 |

#### Top 5 stimuli by accuracy

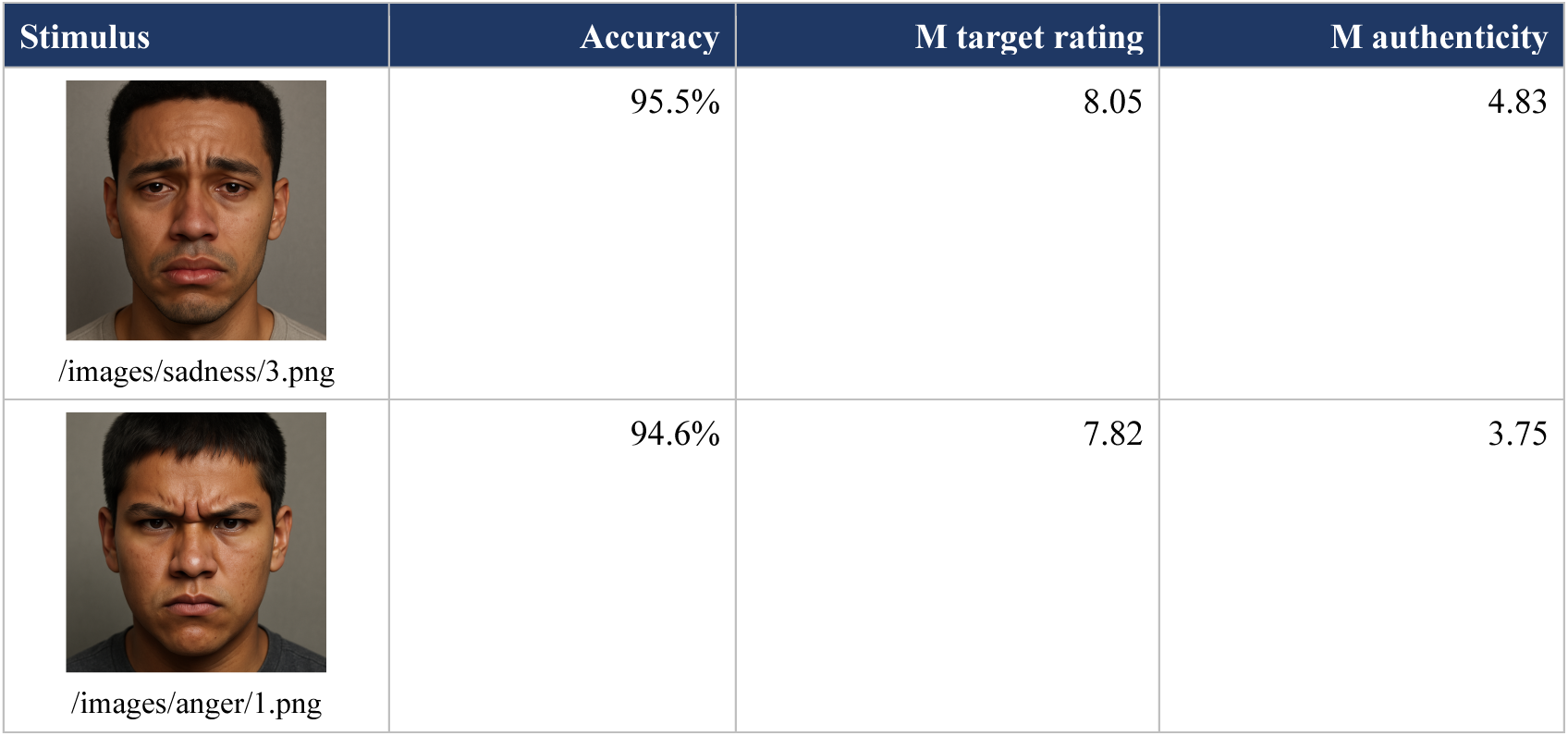
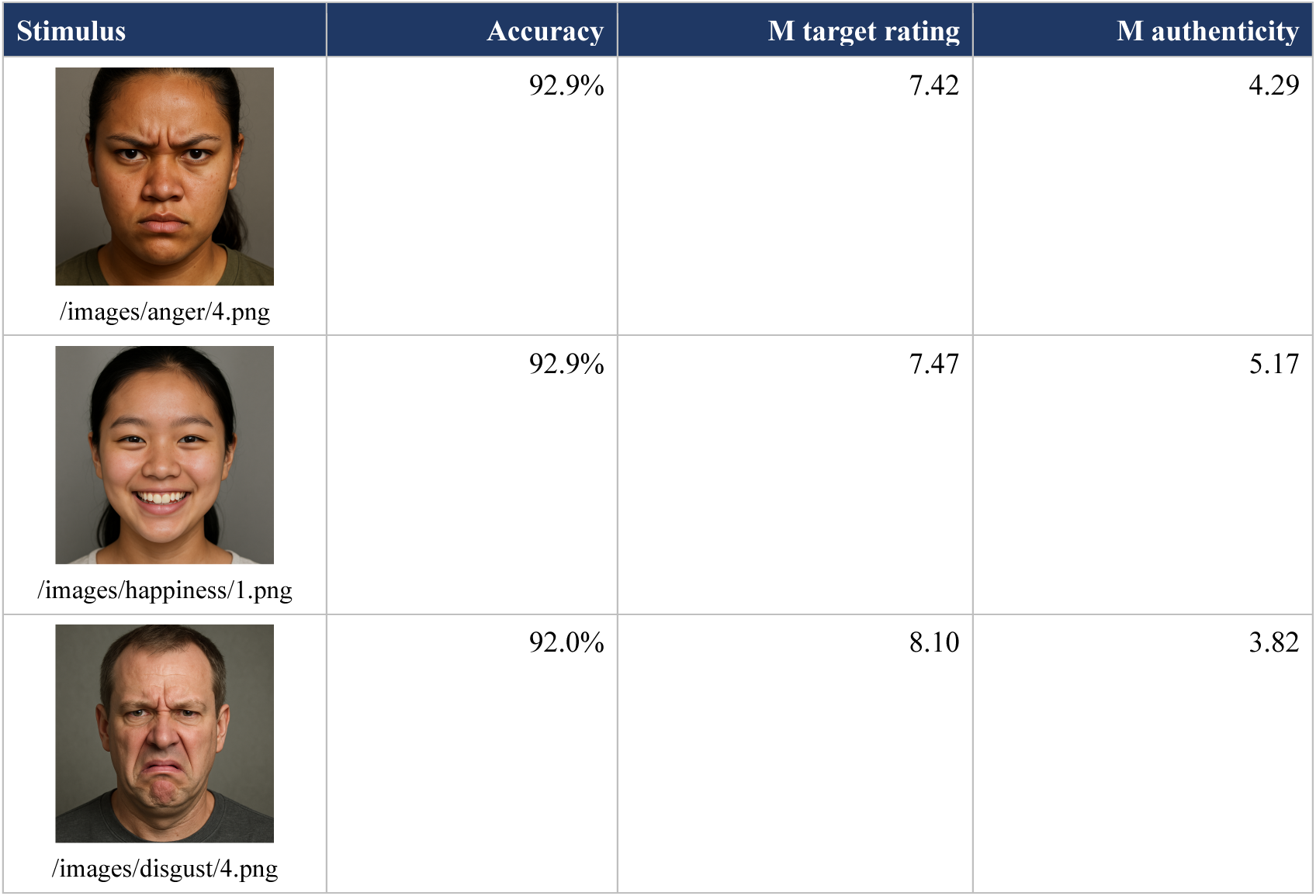

Top 5 stimuli by accuracy.
| Stimulus | Accuracy | M target rating | M authenticity |
| --- | --- | --- | --- |
| /images/sadness/3.png | 95.5% | 8.05 | 4.83 |
| /images/anger/1.png | 94.6% | 7.82 | 3.75 |
| /images/anger/4.png | 92.9% | 7.42 | 4.29 |
| /images/happiness/1.png | 92.9% | 7.47 | 5.17 |
| /images/disgust/4.png | 92.0% | 8.10 | 3.82 |

#### Bottom 5 stimuli by accuracy

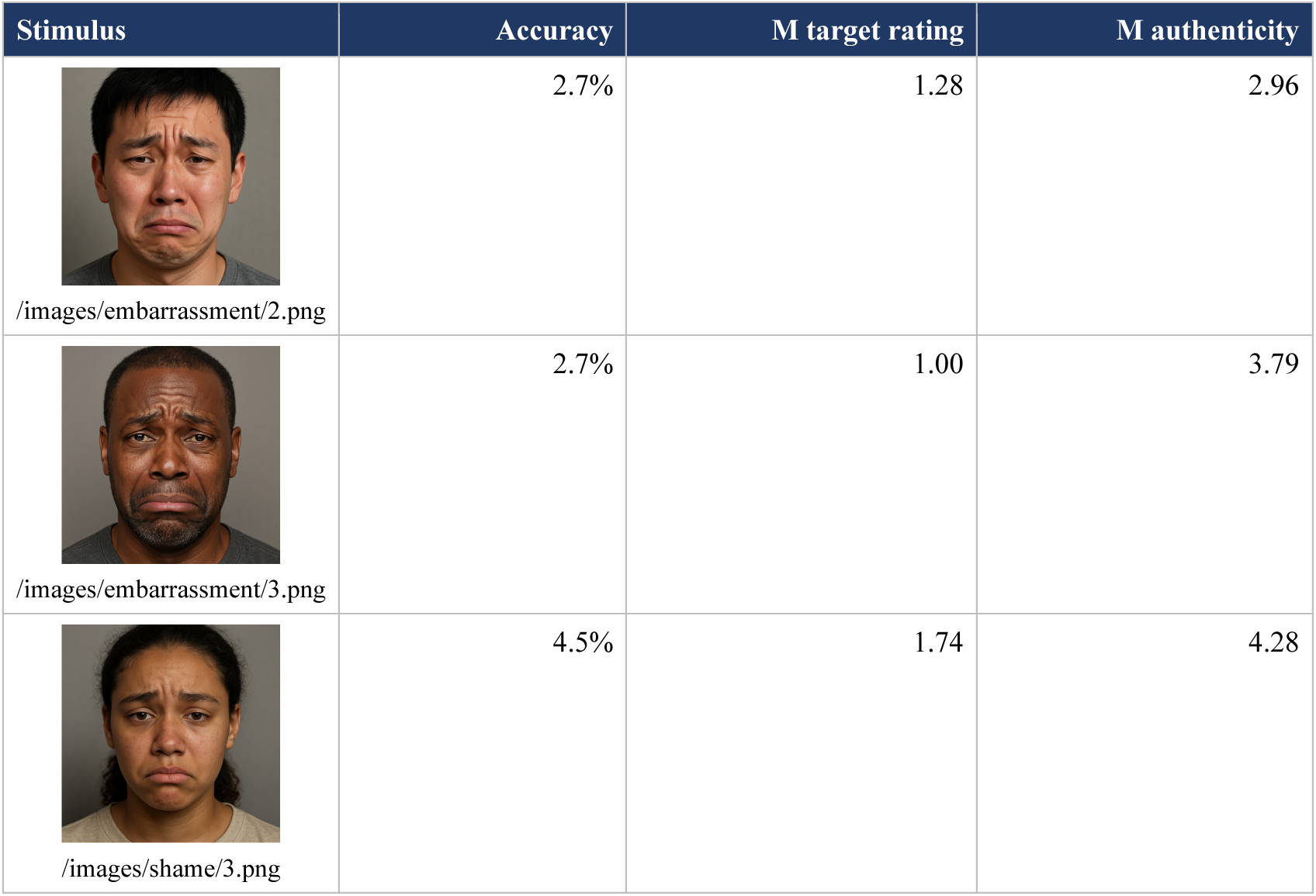
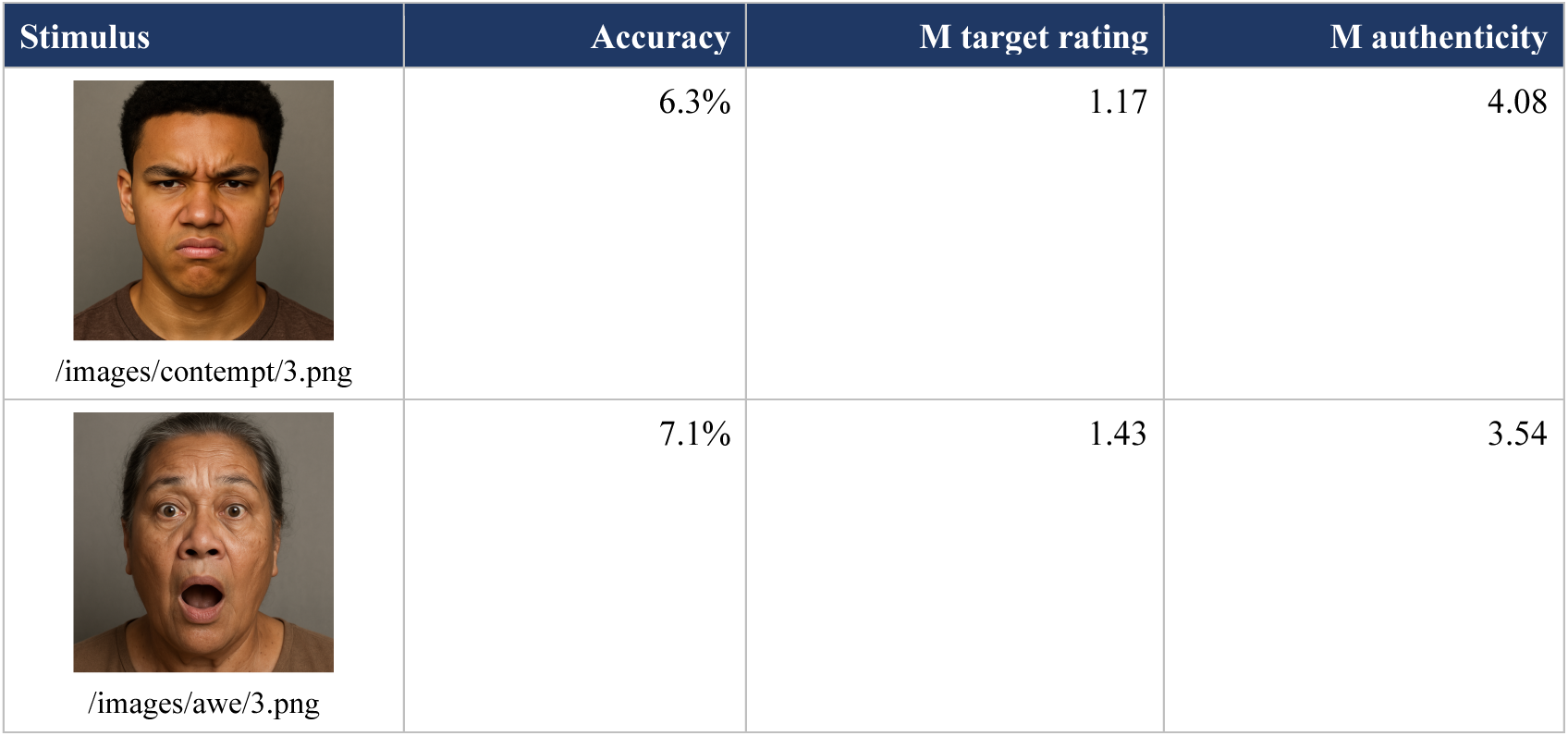

**Bottom 5 stimuli by accuracy.**
| Stimulus | Accuracy | M target rating | M authenticity |
| --- | --- | --- | --- |
| /images/embarrassment/2.png | 2.7% | 1.28 | 2.96 |
| /images/embarrassment/3.png | 2.7% | 1.00 | 3.79 |
| /images/shame/3.png | 4.5% | 1.74 | 4.28 |
| /images/contempt/3.png | 6.3% | 1.17 | 4.08 |
| /images/awe/3.png | 7.1% | 1.43 | 3.54 |

### S6. Verbatim DALL-E 3 prompt template

Every candidate stimulus prompt was instantiated from the following template, with the placeholders *{age}*, *{ethnicity}*, *{gender}*, and *{emotion}* replaced by one combination of values from the orthogonal design described in Methods (Materials, Stimuli):

> “A perfectly centered, ultra-photorealistic front-facing headshot, cropped identically in every image. The top of the subject’s forehead must exactly touch the top edge of the frame, and the bottom of the chin must exactly touch the bottom edge—without any extra space above or below. The subject’s eyes should align horizontally with the midpoint of the image. The subject is a {age} {ethnicity} {gender} displaying {emotion}. The skin has natural texture, realistic lighting, and subtle imperfections for authenticity. The background is a neutral gray. The subject maintains direct eye contact with the camera, and the lighting is soft and even, enhancing facial contours without harsh shadows. The image composition is strictly uniform across all variations.”

### S7. Full demographic distribution of the 52 stimuli

Of the 52 selected stimuli, 26 depicted male subjects and 26 depicted female subjects. Within-emotion gender ratios were 2:2 in seven of the 13 emotions (amusement, anger, contempt, disgust, happiness, interest, pride) and 3:1 in the remaining six (awe, fear, and shame: three female and one male; embarrassment, sadness, and surprise: three male and one female). The seven race/ethnicity categories were represented as follows: Asian, 10; Black, 10; White, 8; American Indian or Alaska Native, 6; Hispanic or Latino, 6; Native Hawaiian or Other Pacific Islander, 6; two or more races, 6. The four age categories were: teenager, 16; young adult, 12; middle-aged adult, 12; older adult, 12.

